# Mutations of SARS-CoV-2 variants of concern escaping Spike-specific T cells

**DOI:** 10.1101/2022.01.20.477163

**Authors:** Nina Le Bert, Anthony Tan, Kamini Kunasegaran, Adeline Chia, Nicole Tan, Qi Chen, Shou Kit Hang, Martin DC Qui, Bianca SW Chan, Jenny GH Low, Barnaby Young, Kee Chong Ng, Derrick Wei Shih Chan, David Chien Lye, Antonio Bertoletti

## Abstract

The amino acid (AA) mutations that characterise the different variants of concern (VOCs), which replaced the ancestral SARS-CoV-2 Wuhan-Hu-1 isolate worldwide, provide biological advantages such as increased infectivity and partial escape from humoral immunity. Here we analysed the impact of these mutations on vaccination- and infection-induced Spike-specific T cells. We confirmed that, in the majority of infected or vaccinated individuals, different mutations present in a single VOC (Delta) or a combined mosaic of more than 30 AA substitutions and deletions found in Alpha, Beta, Gamma, Delta and Omicron VOCs cause modest alteration in the global Spike-specific T cell response. However, distinct numerically dominant Spike-specific CD4 and CD8 T cells preferentially targeted regions affected by AA mutations and do not recognise the mutated peptides. Importantly, some of these mutations, such as N501Y (present in Alpha, Beta, Gamma, and Omicron) and L452R (present in Delta), known to provide biological advantage to SARS-CoV-2 in terms of infectivity also abolished CD8 T cell recognition.

Taken together, our data show that while global mRNA vaccine- and infection-induced Spike-specific T cells largely tolerate the diverse mutations present in VOCs, single Spike-specific T cells might contribute to the natural selection of SARS-CoV-2 variants.

---

Living organisms, including viruses, constantly evolve to adapt to their environment. They acquire random mutations during replication and deleterious or neutral mutations are purged. Mutations that aid to their spread and persistence^1^ and provide a selective advantage become dominant^2,3^.

The current SARS-CoV-2 pandemic illustrates this phenomenon: after an initial year of relative evolutionary stasis, variants have emerged to replace the initial SARS-CoV-2 Wuhan-Hu-1 strain^2,3^. In July 2021 SARS-CoV-2 variants of concern (VOCs) declared by the World Health Organization comprised B.1.1.7-Alpha, B.1.351-Beta, P.1-Gamma, B.1.617.2-Delta^3^. From end-November 2021, a new VOC called Omicron was declared and has spread rapidly worldwide^4^.

The selection of these variants likely occurs due to the combined effect of immunological pressure and acquisition of advantages in transmissibility and fitness^1,5^. Amino acid (AA) variations occur throughout the entire genome^2,3^, however, the most-studied mutations are in the Spike protein^6–11^. Some mutations increase binding to the ACE-2 receptor^12–14^, while others reduce antibody neutralization^6,9,11,15^.

In contrast, the role of SARS-CoV-2-specific T cells in selection pressure has been largely discounted, despite scattered observations that mutations can affect T cell recognition^9,12,16,17^. The main argument is that T cells recognise various epitopes in vaccinated and convalescent individuals^7,18,19^. As such, VOC-mutations are unlikely to alter all of them^7,10^.

This argument is supported by recent reports documenting the ability of Spike-specific T cells to tolerate the high number of mutations present in Omicron^20–26^, and likely the immunological basis of why vaccinations are still highly effective against VOCs in reducing severe COVID-19^27^.

However, the role for multi-specific antiviral immunity is not involvement in the selection process of VOCs should not be entirely discounted. Both, humoral and cellular immunity are multi-specific^28,29^ and while most VOCs fully escape recognition of monoclonal antibodies^6,30^, their ability to escape serum neutralisation from convalescent or vaccinated individuals was less dramatic^6,15^ before the surge of Omicron. In addition, a hierarchy of antiviral efficacy exists within the polyclonal cellular immune response^29^, and thus some mutations might escape the dominant antiviral T cells, similarly to what has been observed for polyclonal antibodies^15^.

## Results

### Breadth of the Spike-specific T cell response

To understand whether Spike-specific T cells uniformly recognise different Spike regions^29^, we designed seven pools of 33-39 overlapping 15-mer peptides covering 180-200 AA long regions (Fig S1A, Table S1). Peptide-reactive cells were quantified by IFN-γ ELISpot ex vivo in 35 vaccinated and 31 convalescents individuals. The mean quantity was different (Fig. S1B, S1C), likely reflecting the time of measurement since T cell induction (3 versus 12 months, respectively). Yet, important commonalities were detected. First, as already described^31^, we found significant heterogeneity in the quantity of Spike-specific T cells (Fig. S1B, S1C). Second, most of the individuals exhibited T cells recognising all seven distinct peptide pools (Fig. 1A), in line with the reported T cell multi-specificity^7,18,19^. However, a dominant T cell response towards a single peptide pool was frequently observed. In 8/35 vaccinated and 9/31 convalescents, with T cells specific for a single pool exceeding 40% of the total Spike-specific T cell response (Fig. 1B). The Spike region 886-1085 was the most immunogenic in both vaccinated (65%) and convalescents (34%) (Fig. 1C). This region is fairly conserved among different VOCs, but includes the D950N, S982A and T1027I mutations in the Delta, Alpha and Gamma VOCs, respectively, and the mutations S954H, S969K, S981F in Omicron. In 16% of vaccinated and 19% of convalescents, a dominant T cell response was observed towards the region 336-510 (Fig. 1C), containing several VOC mutations that affect the receptor binding affinity (N501Y^13,14^, L452R^12^), antibody (K417^32^, T478K^15^, E484K^15^) and T cells (L452R^12^) recognitions.

**Figure 1:**
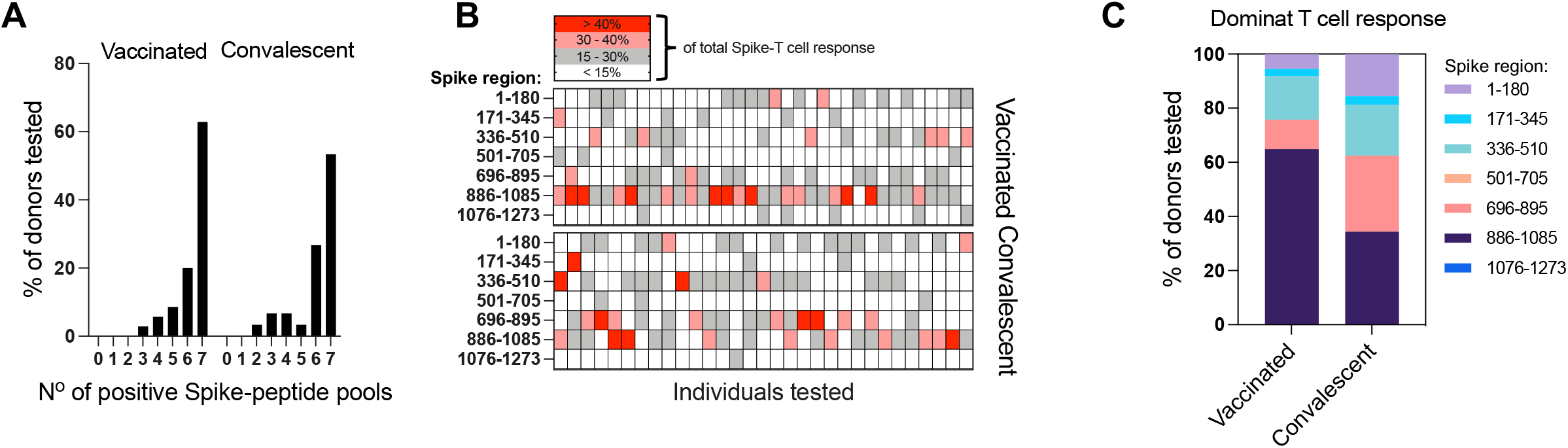
Breath of the Spike-specific T cell response in BNT162b mRNA vaccinated donors and SARS-CoV-2 convalescents. **A**, Bar graphs show the percentage of donors (vaccinated n=35; convalescent n=31) reacting to the number of Spike-peptide pools tested (total 7 distinct peptide pools). **B**, Heatmap is indicating the percentage of the response towards a single peptide pool in proportion to the total Spike-specific response in each of the tested individuals. **C**, Percentage of tested individuals with a dominant response to one of the 7 peptide pools is shown.

### Impact of VOC mutations on global Spike-specific T cells

Next, we tested the effect of the mutations on Spike-specific T cells. First, we aimed to understand in 100 individuals the impact of Delta mutations on Spike-specific T cells induced by BNT162b2 vaccine. We stimulated whole blood with three peptide pools covering the whole Spike protein (253 peptides) and the regions mutated in Delta (24 peptides) with and without the AA-substitutions/deletions (Fig 2A, Table S2).

**Figure 2:**
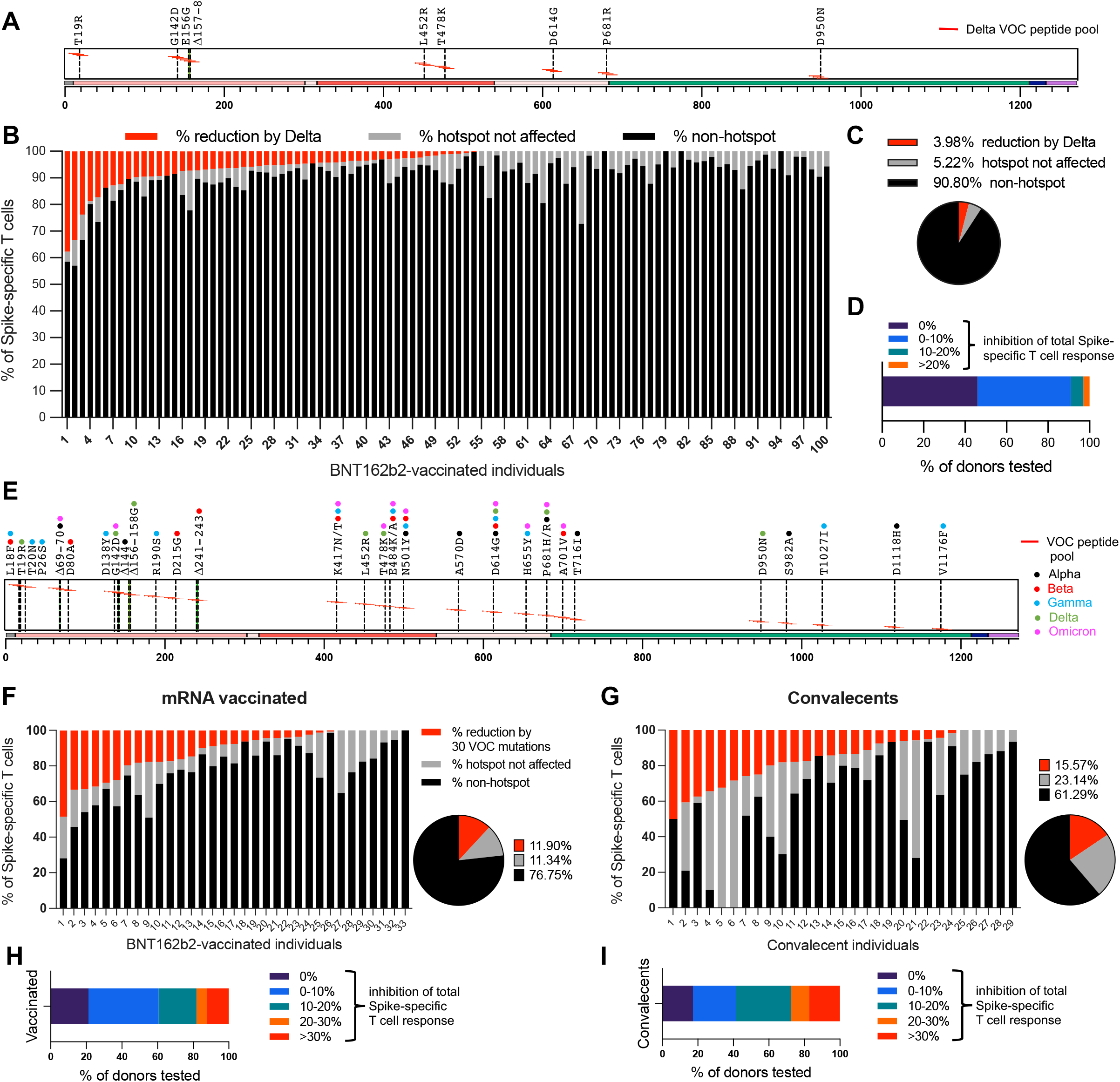
Impact of VOC mutations on the global Spike-specific T cell response. **A**, Schematic of Spike protein with the amino acid (AA) substitutions/deletions characteristic for the Delta VOC. Red lines indicate the locations of the 15-mer peptides covering the mutated regions, which were tested as separate pools with and without the AA substitutions/deletions. **B**, Whole blood of n=100 BNT162b2 vaccinated donors was stimulated with 3 separate peptide pools covering 1) the whole Spike protein, 2) the Delta mutations without and 3) with the AA substitutions/deletions. IFN-γ release was quantified after overnight stimulation in the plasma. Bars indicate the percentage of the Spike-peptide response targeting the conserved regions (black), the mutated regions but not affected (grey) and the percentage inhibited by the mutations (red). **C**, Mean reduction of the response to Delta AA substitutions/deletions in proportion to the total Spike-response. **D**, Proportion of donors whose response was reduced by the indicated percentages. **E**, Schematic of Spike protein with 30 tested AA substitutions/deletions characteristic for the VOCs (Alpha, Beta, Gama, Delta, Omicron). Red lines indicate the locations of the 15-mer peptides covering the mutated regions, which were tested as separate pools with and without the AA substitutions/deletions. PBMCs of 33 BNT162b2 vaccinated donors (**F**) and 29 SARS-CoV-2 convalescents (**G**) was stimulated with 3 separate peptide pools covering 1) the whole Spike protein, 2) the VOC mutations without and 3) with the AA substitutions/deletions. Bars indicate the percentage of the Spike-peptide response targeting the conserved regions (black), the mutated regions but not affected (grey) and the percentage inhibited by the mutations (red). Pie charts are indicating the mean reduction of the response to 30 tested VOC AA substitutions/deletions in proportion to the total Spike-response. Proportion of vaccinated donors (**H**) and convalescents (**I**) whose response was reduced by the indicated percentages.

We measured the magnitude of the Spike-specific T cell response, its proportion targeting peptides affected by AA changes and calculated the reduction when stimulated with peptides containing Delta-specific mutations (Fig. 2B), which was on average 3.9% (Fig. 2C). The Spike-specific T cell response was not affected in 46% of the vaccinated individuals; 45% showed less than 10% reduction and only in 3% did we observe more than 20% reduction in Spike-specific T cells (Fig. 2D). Of note, we confirmed in this large population that there is vast heterogeneity of the Spike-specific T cell response in different individuals (Fig. S2).

Second, we defined the combined impact of the mutations that are characteristic of the Alpha, Beta, Gamma and Delta VOCs on Spike-specific T cells in 33 vaccinated and 29 convalescents (Fig. 2E, Table S2). Note, 10 of these mutations are also characteristic of the newly emergent Omicron VOC^4^ (Table S3). The combined effect of 30 AA mutations was tested with IFN-γ ELISpot assays.

This combination of 30 AA mutations reduced the T cell response on average by 11.9% (vaccinated, Fig 2F) and 15.6% (convalescent, Fig 2G), a greater reduction than that detected for the Delta mutations alone (3.9%). Spike-specific T cells were not affected in 7/33 (21%) mRNA vaccinated (Fig. 2H) and 5/29 (17%) convalescent individuals (Fig. 2I). Less than 10% reduction was observed in 39% of vaccinated and 24% of convalescents. Only in 1 individual of both groups did the combined VOC mutations reduce the Spike-specific T cells by 50%.

### Definition of single-peptide specificities of dominant and subdominant Spike-specific T cells

Next, we characterized epitope-specificity and CD4/CD8 phenotype of vaccine- and infection-induced T cells. We utilized an unbiased approach, based on PBMC stimulation with a single peptide pool covering whole Spike and expansion of specific T cells. The T cell lines were then used to confirm the single-peptide specificity, define the phenotype of the responsive T cell (CD4/CD8) and, in selected cases, their HLA-Class I restriction (Fig. S3–5). This approach allowed us to define the dominant T cell specificities, irrespective of the HLA-Class I and Class II profile of the tested individuals.

First, we showed that the expansion procedure preserved the overall hierarchy of Spike-specific T cell recognition detected ex vivo in most of the tested individuals and it was remarkably stable across different time points (Fig. 3SB).

**Figure 3:**
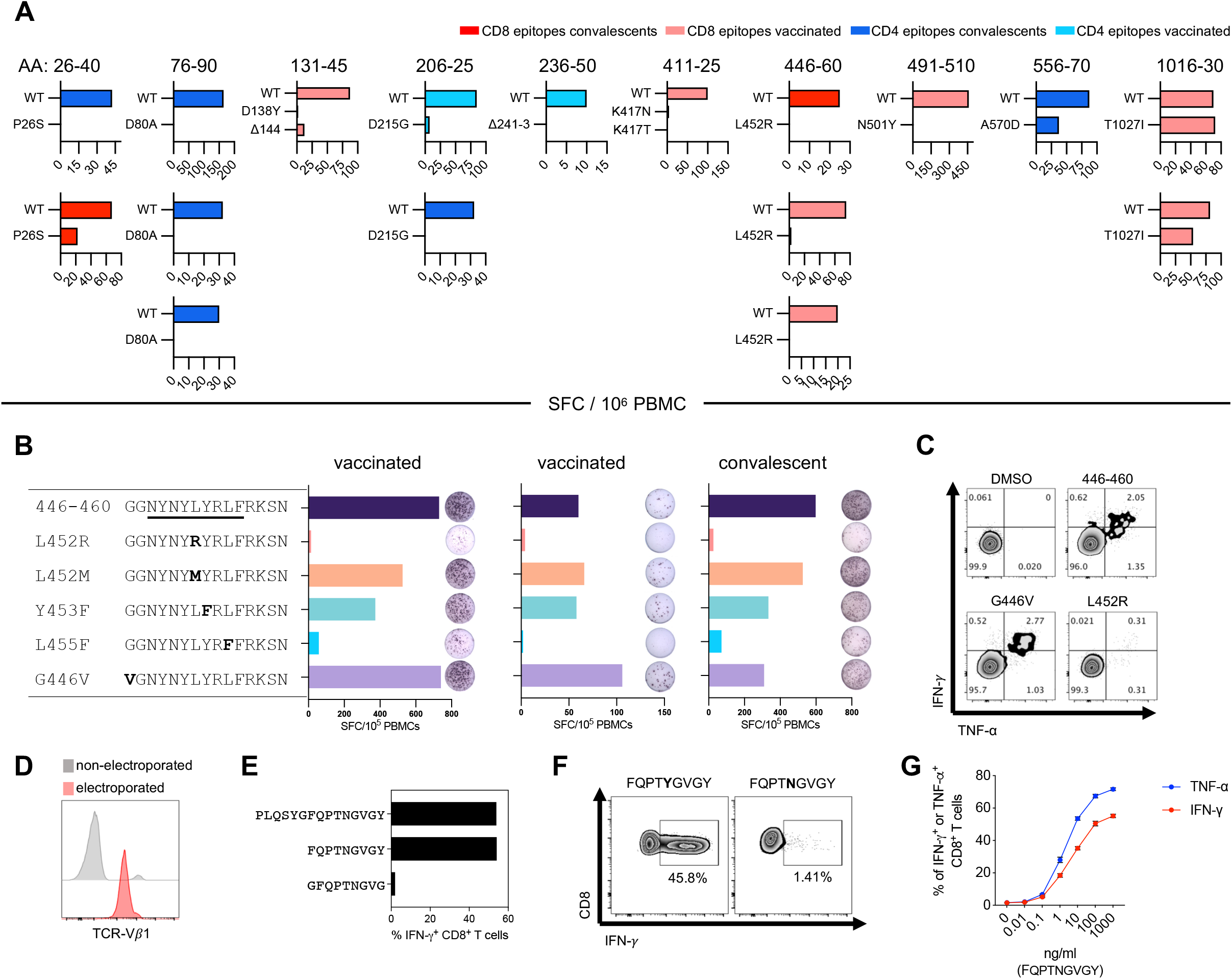
Impact of VOC AA mutations on Spike-specific T cells. **A**, PBMCs of the individuals in whom the T cell specificities were defined (and cells were available) were stimulated ex vivo in parallel with peptides containing either the wildtype or the mutated sequence. Peptide reactivity was analysed by IFN-γ ELISpot assay. Frequency of spot forming cells (SFC)/106 PBMC are shown. **B**, PBMCs of two vaccinated and one convalescent HLA-A*24:02+ donor were stimulated with the 446-460 peptide for 10 days and subsequently tested in parallel with peptides containing either the wildtype or the five indicated mutated sequences. Peptide response was measured by IFN-γ ELISpot assay. **C**, The 446-460 short-term T cell line of a vaccinated HLA-A*24:02+ donor was stimulated in parallel with peptides containing either the wildtype sequence (446-460), or the G446V and L452R mutations. Peptide response was measured by intracellular cytokine staining. **D**, TCR Vβ1 antibody was used to detect the introduced TCR. Representative FACS histogram plots showing TCR Vβ1 expression in allogenic T cells 24 hours after mRNA electroporation. **E**, TCR-redirected T cells stimulated in parallel with the 15-mer peptide 491-505 (PLQSYGFQPTNGVGY), 9-mer peptides 497-505 (FQPTNGVGY) and 496-504 (GFQPTNGVG) analysed by intracellular cytokine staining. **F**, TCR-redirected T cells stimulated in parallel with peptide 497-505 with and without the N501Y mutation. Peptide reactivity was analysed by intracellular cytokine staining. **G**, TCR-redirected T cells stimulated with a 10-fold dilution series of the 497-505 (FQPTNGVGY) peptide and analysed by intracellular cytokine staining.

Subsequently, we defined the single peptide specificity and the CD4/CD8 phenotype of T cells in nine vaccinated (Fig. S3C) and 11 convalescents (Fig. S4). Eighteen distinctive Spike-specific CD4 epitopes and 17 different CD8 epitopes were characterized (Fig. S3C, S4; Table1). HLA-Class I restriction of 6 distinct CD8 T cell lines was identified utilizing EBV-B cell lines with shared/non-shared HLA-Class I molecules (Fig. S5).

Surprisingly, among the defined peptides containing epitopes targeted by Spike-specific T cells, 12/18 CD4 T cells and 10/17 CD8 T cells recognized peptides that contain 30 out of the 55 distinct AA mutations characteristic of the five VOCs (Table 1, Table S2).

**Table 1:**
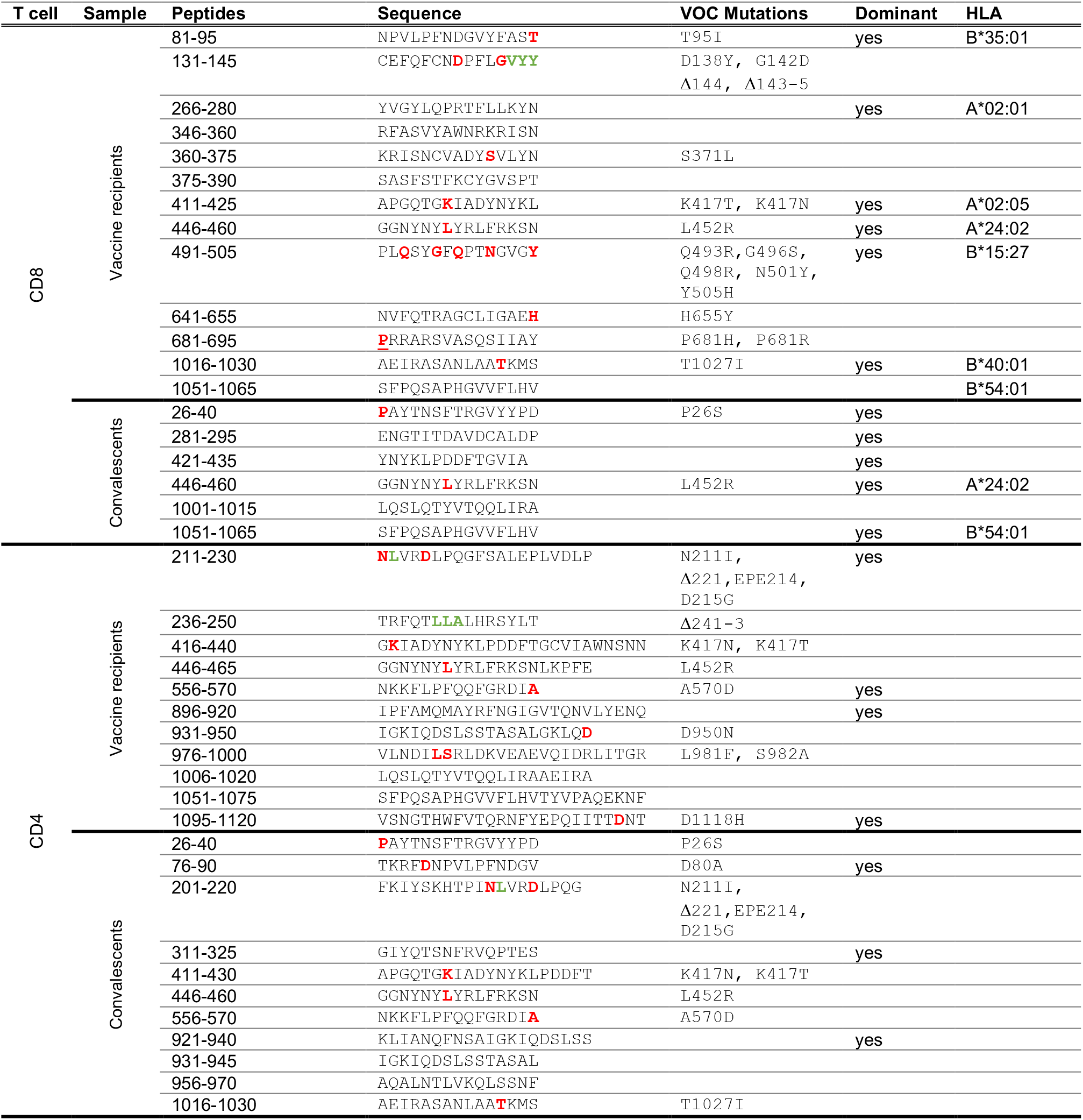
Details of identified Spike-specific T cell specificities

### Mutations affecting Spike-specific T cells

The impact of the AA mutations affecting the peptide-specific T cells was analysed both directly ex vivo (Fig. 3A) and in T cell lines (Fig. S6). We tested 6 CD8 and 5 CD4 T cell specificities containing AA mutations. PBMCs of the individuals in whom the T cell specificities were defined were stimulated in parallel with peptides containing either the wildtype or the mutated AA sequence.

All AA mutations affected the T cell recognition with the partial exception of mutation T1027I present in S1016-30. This mutation altered T cell recognition in one out of two individuals tested. HLA-restriction (Fig. S5) and visualization with HLA-pentamer on the individual who tolerated T1027I (Fig. S7) showed that the T cells were B40-restricted and recognized the epitope S1016-1024, which lay outside of the mutations. The ability of the other AA mutations to inhibit T cell recognition strongly suggest that they were located within the T cell epitopes. Mutations affected both CD4 (i.e. S76-90, S206-25, S236-50) and CD8 T cell responses (i.e. S411-25, S446-60, S491-510). Of note, peptide S26-40 stimulated a CD4 and a CD8 T cell response in two different convalescents, suggesting this peptide contains two distinct epitopes. P26S strongly inhibited both CD4 and CD8 T cells.

Of particular interest, we demonstrated that HLA-A*02:05-restricted CD8 T cells specific for the peptide 411-25 were completely inhibited by Beta and Omicron mutation K417N and by Gamma mutation K417T. Moreover, we observed that the mutations L452R and N501Y, which have negligible effect on antibody recognition ^33,34^ but increase the infectivity of the Delta^12^ and Beta/Gamma^13,14^ VOCs respectively, clearly inhibited recognition of CD8 T cells specific for the HLA-A*24:02 epitope GNYNYLYRLF and HLA-B*15:27 epitope FQPTNGVGY.

### Impact of L452R and N501Y on Spike-specific T cells

The observation that L452R and N501Y, present in Delta (L452R), Alpha, Beta, Gamma and Omicron (N501Y), respectively, increase infectivity^12^ and concomitantly abolish CD8 T cell recognition prompted us to analyse these CD8 T cells in more detail.

The region S446-60 contains the HLA-A*24:02 epitope 448-56 (NYNYLYRLF), has been shown by Motozono et al^12^ to induce a dominant CD8 T cell response in convalescents and to be inhibited by L452R. Here we demonstrated that S448-56-specific CD8 T cells are also elicited in A*24:02+ vaccinated individuals and we confirmed the HLA-A*24:02 restriction (Fig. S5).

We then tested the ability of L452R to inhibit T cell activation in comparison with other reported AA mutations that affect the Spike region 446-60, namely L452M, Y453F, L455F and G446V (Fig. 3B,C). The mutation Y453F was characteristic of the SARS-CoV-2 strain infecting minks^35^, while L452M, Y453F, L455F and G446V, have been occasionally detected^36^. CD8 T cell lines specific for Spike 446-60 were generated in three different HLA-A*24:02+ individuals and we tested the impact of the 5 distinctive AA substitutions. L452R abolished the CD8 T cell recognition almost completely in all the three individuals, followed by L455F, while the other AA substitutions (G446V, L452M, Y453F) were better tolerated (Fig. 3B,C).

To characterize the HLA-B*15:27 restricted CD8 T cell response to the Spike region containing the mutation N501Y, we engineered T cell receptor (TCR)-redirected T cells. The alpha and beta TCR chains of T cells activated by peptide 491-505 were sequenced and cloned into a pVAX1 vector, which allowed the expression of the introduced TCR in allogenic PBMC (Fig 3D). The TCR-redirected T cells were used to define the short epitope (FQPTNGVGY; Fig. 3E) and we confirmed the ability of N501Y to completely abolish CD8 T cell recognition (Fig. 3F). Titration of the peptide concentration used for T cell activation demonstrated the high affinity of the TCR, a concentration as low as 0.1 ng/ml induced a response which plateaued at 1 μg/ml (Fig. 3G).

## Discussion

We designed experiments to address two questions: whether VOCs can escape the global Spike-specific T cell response induced by infection or vaccination and whether T cells can play a role in VOCs selection.

We confirmed the marked multi-specificity of Spike-specific T cells. Although we observed a hierarchy among the T cells recognizing different Spike regions, the dominant Spike-specific T cells rarely occupied more than 40% of the repertoire. Furthermore, the region 886-1085 that is preferentially targeted by T cells contains few mutations present in Alpha (S982A), Gamma (T1027) and Delta (D950N) ^15^ and also in the newly emerged Omicron VOC (S954H, S969K, S981F)^4^.

The broad multi-specificity translated into the functional ability to largely tolerate AA substitution present in different VOCs. This was first observed in a large population of mRNA vaccinated individuals (n=100). In 91% of them, the effect of mutations present in Delta, inhibited Spike-specific T cells by less than 10%. Second, when we tested the combined effect of 30 distinct AA substitutions/deletions (mutations found in Alpha, Beta, Gamma and Delta VOCs, of which 10 have also been detected in Omicron), we found that convalescent or vaccinated individuals of Asian origin possess Spike-specific T cells that largely tolerate the combined AA substitutions. These data agree with the recent reports of the substantial, although not absolute, preservation of global Spike-specific T cell response against the highly mutated Omicron^20–26^.

Our data however reveals novel important features of the Spike-specific cellular immunity. In addition to the detection of a broad heterogeneity in the magnitude of the T cell response, we demonstrated that 12 distinct AA substitutions of VOCs alter 11 individual T cell specificities characterized in a relatively small number of vaccinated and convalescent individuals (n=20). VOC mutations affect the activation of CD8 and CD4 T cells and mutations already known to abrogate antibody recognition like K417N^34^ (present in Beta and Omicron VOC) or K417T and D138Y (present in Gamma VOC)^8^ also inhibit Spike-specific CD8 T cells.

Remarkably, we observed that mutations L452R (present in Delta) and N501Y (present in Alpha, Beta Gamma and Omicron), which have negligible effect on the neutralization ability of polyclonal sera^33,34^, but increase the binding affinity of Spike protein to the ACE2 receptor^12–14^, abolish the recognition of CD8 T cells specific for one HLA-A*24:02 (S447-56) and one HLA-B*15:27 (S497-505)-restricted epitope. This convergence of biological effects is reminiscent of the hypothesis of the causes of influenza hemagglutinin variant selection. In animal models of Influenza, antibodies select viruses with mutations that provide a generalized advantage by increasing receptor avidity^5^.

In general, immune escape mutations affecting T cell epitopes are selected in individuals with chronic viral infections (i.e. HIV, HBV, HCV, HDV, HCMV, EBV)^37^ but are unlikely to uniformly affect the wider human population. Since the biology of T cell recognition makes the T cell repertoire largely unique for each individual^29^, it is unlikely that a virus variant with a single set of immune escape mutations will affect the whole human population identically and successfully spread globally. However, if mutations permit escape from a specific T cell specificity in parallel with increasing the receptor binding affinity of Spike, as in the case of L452R and N501Y, such advantage will no longer be solely restricted to a selected population. Interestingly, recent mathematical models of SARS-CoV-2 variant spread suggested that mutations able to concurrently increase infectivity and immune escape are likely to rapidly propagate in the population^38^.

Our data do not provide the demonstration that this chain of events took place during this pandemic. Nevertheless, they indicate an alternative possibility to the prevalent theory that postulates that VOCs emerged exclusively under the pressure of neutralizing antibodies.

We made the unexpected observation that a large number of T cells (20 out of 35) selected in vitro by PBMC stimulation with whole Spike peptide pool recognized peptides carrying the VOC mutated regions of Spike. This is at odds with ex-vivo results obtained by us and others, since T cells specific for variant regions represent a minority of the global Spike T cell repertoire in the ex vivo analysis^7,18–26^. One might speculate that the in vitro expansion mimics the immunological events occurring after SARS-CoV-2 infection and, as such, the analysis of T cells after in vitro expansion selects for the dominant effector T cell response present during the acute phase of response. These dominant T cells might thus exert higher selective pressure. Future studies will be necessary to verify the robustness of the observation and its real mechanisms. However, by determining HLA-restrictions of a number of these T cell epitopes targeting mutated regions (at least for HLA-Class I), we provide the possibility to test whether, for example, breakthrough infection might occur more frequently in individuals with such HLA-Class I profiles. Finally, the demonstration that AA mutations escaping specific CD8 T cells and concomitantly offer the virus a biological advantage in term of increased infectivity^14^, provides the theoretical possibility that T cell pressure might contribute to the selection of VOCs.

There are some limitations to this study. We tested the impact of AA substitutions on Spike-specific T cells utilizing peptide-pulsed target cells and not infected cells. This method might overestimate the quantity of T cells specific for SARS-CoV-2 because low affinity T cells might have been quantified and be more sensitive to AA mutations. On the other hand, peptide-pulsed target cells cannot be used to evaluate the impact that AA substitutions might have on the processing of T cell epitopes^37^. AA outside the T cell epitopes can alter their generation^39^ and as such we might have underestimate the impact of the mutations on the Spike-specific cellular immunity.

## Methods

### Ethics statement

All donors provided written consent. The study was conducted in accordance with the Declaration of Helsinki and approved by the NUS institutional review board (H-20-006) and the SingHealth Centralised Institutional Review Board (reference CIRB/F/2018/2387).

### Human samples

Donors were recruited based on their clinical history of SARS-CoV-2 infection and their vaccination status. Blood samples of recovered COVID-19 patients (n=35) were obtained 6-12 months post PCR negativity. Blood samples of two vaccinated cohorts were taken, 35 donors were recruited at multiple timepoints until 3 months post second dose BNT162b, and additional 100 donors at 9 months post second dose BNT162b. Vaccinated individuals were all healthy adults, 20-65 years old, of Asian origin.

### PBMC isolation

Peripheral blood mononuclear cells (PBMC) were isolated by density-gradient centrifugation using Ficoll-Paque. Isolated PBMC were either studied directly or cryopreserved and stored in liquid nitrogen until used in the assays.

### Peptide pools

15-mer peptides overlapping by 10 amino acids spanning the entire protein sequence of SARS-CoV-2 Spike were synthesized (GenScript; see Table S1). To stimulate whole blood or PBMC, the peptides were divided into 7 pools of about 40 peptides. For single peptide identification, peptides were organized in a matrix of 16 numeric and 16 alphabetic pools. Peptides with and without VOC mutations were mixed into two separate pools (Table S2).

### Cytokine release assay (CRA) from whole peripheral blood

320 μl of whole blood drawn on the same day were mixed with 80 μl RPMI and stimulated with the indicated SARS-CoV-2 Spike peptide pools at 2 μg/ml or with DMSO as a control. After 16 hours of culture, the culture supernatant (plasma) was collected and stored at −80°C. Cytokine concentrations in the plasma were quantified using an Ella machine with microfluidic multiplex cartridges measuring IFN-γ and IL-2 following the manufacturer’s instructions (ProteinSimple). The level of cytokines present in the plasma of DMSO controls was subtracted from the corresponding peptide pool stimulated samples. The positivity threshold was set at 10x times the lower limit of quantification of each cytokine (IFN-γ = 1.7pg/ml; IL-2 = 5.4pg/ml) after DMSO background subtraction.

### ELISpot assay

ELISpot plates (Millipore) were coated with human IFN-γ antibody (1-D1K, Mabtech; 5 μg/ml) overnight at 4°C. 400,000 PBMC were seeded per well and stimulated for 18h with pools of SARS-CoV-1/2 peptides (2 μg/ml). For stimulation with peptide matrix pools or single peptides, a concentration of 5 μg/ml was used. Subsequently, the plates were developed with human biotinylated IFN-γ detection antibody (7-B6-1, Mabtech; 1:2000), followed by incubation with Streptavidin-AP (Mabtech) and KPL BCIP/NBT Phosphatase Substrate (SeraCare). Spot forming units (SFU) were quantified with ImmunoSpot. To quantify positive peptide-specific responses, 2x mean spots of the unstimulated wells were subtracted from the peptide-stimulated wells, and the results expressed as SFU/106 PBMC. We excluded the results if negative control wells had >30 SFU/106 PBMC or positive control wells (PMA/Ionomycin) were negative.

### Flow Cytometry

PBMC or expanded T cell lines were stimulated for 5h at 37°C with or without SARS-CoV-2 peptides (2 μg/ml) in the presence of 10 μg/ml brefeldin A (Sigma-Aldrich). Cells were stained with the yellow LIVE/DEAD fixable dead cell stain kit (Invitrogen) and anti-CD3 (clone SK7; 3:50), anti-CD4 (clone SK3; 3:50), and anti-CD8 (clone SK1; 3:50) antibodies. Cells were subsequently fixed and permeabilized using the Cytofix/Cytoperm kit (BD Biosciences-Pharmingen) and stained with anti-IFN-γ (clone 25723, R&D Systems; 1:25) and anti-TNF-α (clone MAb11; 1:25) antibodies and analyzed on a BD-LSR II FACS Scan. Data were analyzed by FlowJo (Tree Star Inc.). Antibodies were purchased from BD Biosciences-Pharmingen unless otherwise stated.

### Expanded T cell lines

T cell lines were generated as follows: 20% of PBMC were pulsed with 10 μg/ml of the overlapping SARS-CoV-2 peptides (all pools combined) or single peptides for 1 hour at 37°C, subsequently washed, and cocultured with the remaining cells in AIM-V medium (Gibco; Thermo Fisher Scientific) supplemented with 2% AB human serum (Gibco; Thermo Fisher Scientific). T cell lines were cultured for 10 days in the presence of 20 U/ml of recombinant IL-2 (R&D Systems).

### HLA-restriction assay

The HLA-haplotype (4 digit HLA-typing) of individuals was determined and different EBV transformed B cells lines with one common allele each were selected for presentation of the indicated peptides. B cells were pulsed with 10 μg/ml of the peptide for 1 hour at 37°C, washed three times, and cocultured with the expanded T cell line at a ratio of 1:1 in the presence of 10 μg/ml brefeldin A (Sigma-Aldrich). Non-pulsed B cell lines served as a negative control detecting potential allogeneic responses and autologous peptide-pulsed cells served as a positive control.

### TCR-redirected SARS-CoV-2-specific CD8+ T cells

Spike-specific T cell line was stimulated for 5 hours with peptide S491-505 and the activated antigen-specific T cells were identified through the expression of CD107a. The CD107a+ T cells were sorted and single cell TCR sequencing was performed and analyzed using the 10x Genomics human T cell V(D)J amplification kit (10x Genomics) according to the manufacturer’s recommendations.

Spike 491-510-specific TCR α and β chain genes were subcloned into T7 expression vector (p-VAX1), the SARS-CoV-2-TCR mRNA was transcribed in vitro using the mMESSAGE mMACHINE™ T7 ULTRA Transcription Kit (ThermoFisher Scientitic) following the manufacturer’s protocols. To introduce the TCR expression in non-memory T cells, PBMCs from healthy individuals were isolated and expanded in vitro for 7 days in the presence of 50 ng/ml of OKT-3 (Miltenyi) and 600 IU/ml IL-2 (R&D Systems) in AIM-V (Gibco) medium supplemented with 2% human AB serum (Gibco). The concentration of IL-2 was increased to 1000 IU/ml on day 7 and the expanded T cells were electroporated with the indicated mRNA at a concentration of 2 pg mRNA/cell on day 8 using 4D Nucleofector™ System (Lonza) according to the manufacturer’s instructions. Electroporated T cells were rested for 5 mins before been maintained in AIM-V media supplemented with 10% human AB serum and 100 IU/ml IL-2 overnight. The expression of the introduced TCR were examined using Live/Dead Fixable Yellow Dead Cell Stain Kit (ThermoFisher Scientific), anti-human TCR Vβ1 (Beckman Coulter), anti-human CD3 and CD8 antibodies (BD Bioscience).

### Peptide-pulse experiment

The EBV-transformed lymphoblastoid B (EBV-B) cell lines with identified HLA-B*15:27 phenotype were used as antigen-presenting cells and cocultured with the indicated concentrations of peptides for 1 hour at 37 °C. The peptide-pulsed EBV-B cells were washed twice with HBSS (Gibco) before cocultured with the TCR-redirected T cells at a 1:1 ratio in the presence of brefeldin A (2 μg/ml) overnight. The cells were stained with Live/Dead in 1 × PBS for 10 min at room temperature and then stained with anti-human CD3 and CD8 antibodies for 30 min at 4°C. The cells were fixed and permeabilized using the Cytofix/Cytoperm fixation/permeabilization (BD Biosciences) buffer following the manufacturer’s protocols. Intracellular cytokine staining was performed with anti-human IFNγ and TNFα (BD Biosciences) antibodies for 30 min at room temperature, followed by washing and analysis by flow cytometry.

---

**Figure S1:**
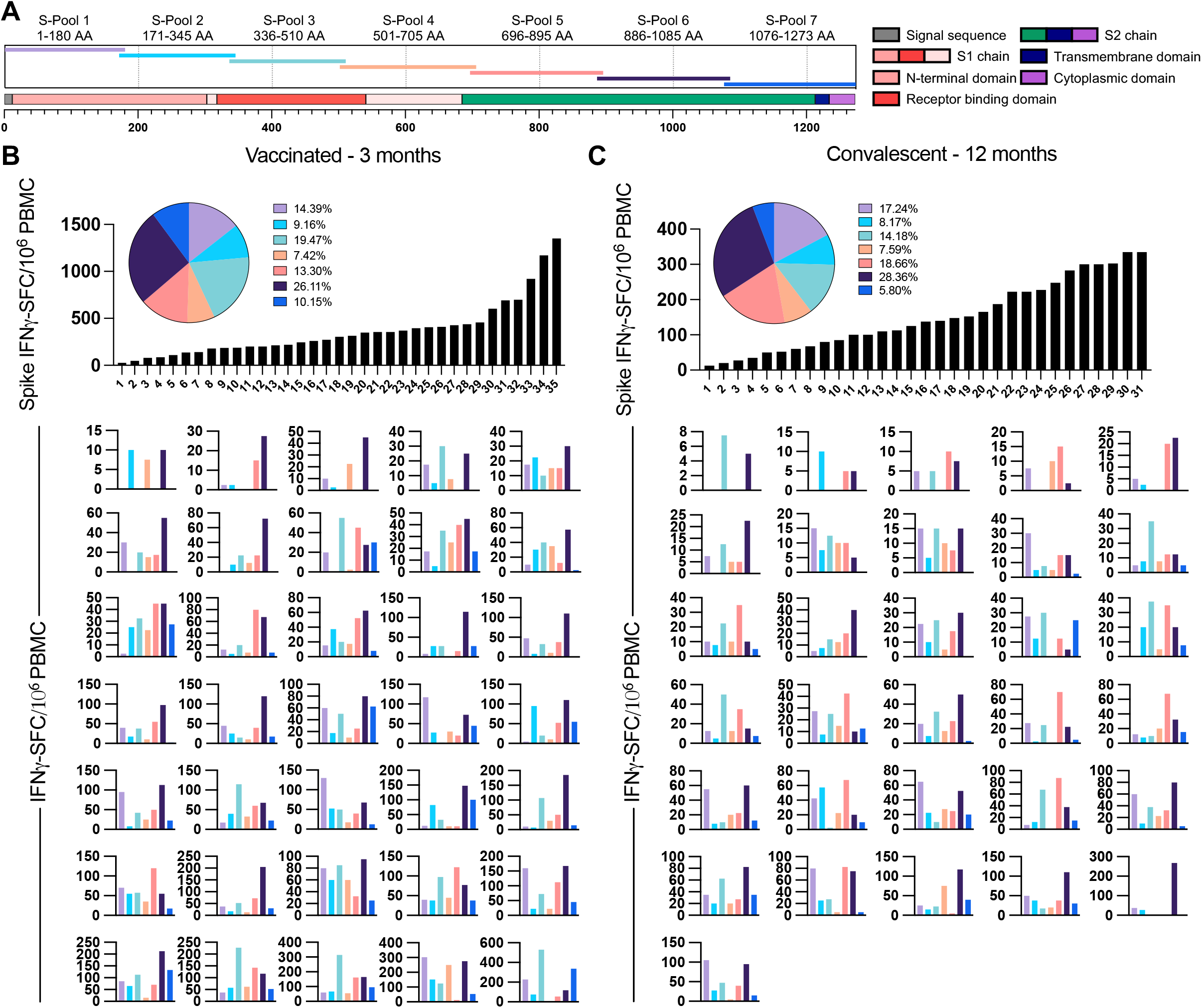
Breath of the Spike-specific T cell response in BNT162b mRNA vaccinated donors and SARS-CoV-2 convalescents. **A**, Schematic of the 7 Spike-specific peptide pools containing 15-mer overlapping peptides spanning the entire Spike protein. **B**, Frequency of peptide-reactive cells in 35 individuals 3 months post 2-dose BNT162b2 vaccination and **C**, frequency of peptide-reactive cells in 31 individuals 12 months post infection with SARS-CoV-2. Black bar graphs in B and C show the total of IFN-γ spot forming cells (SFC)/106 PBMC to all 7 peptide pools combined. The pie charts show the mean proportion of the response to the 7 distinct Spike-peptide pools. Coloured bar graphs below show the frequency of SFC to the 7 distinct Spike-peptide pools in the 35 different vaccinated individuals (B) and in the 31 different convalescents (C).

**Figure S2:**
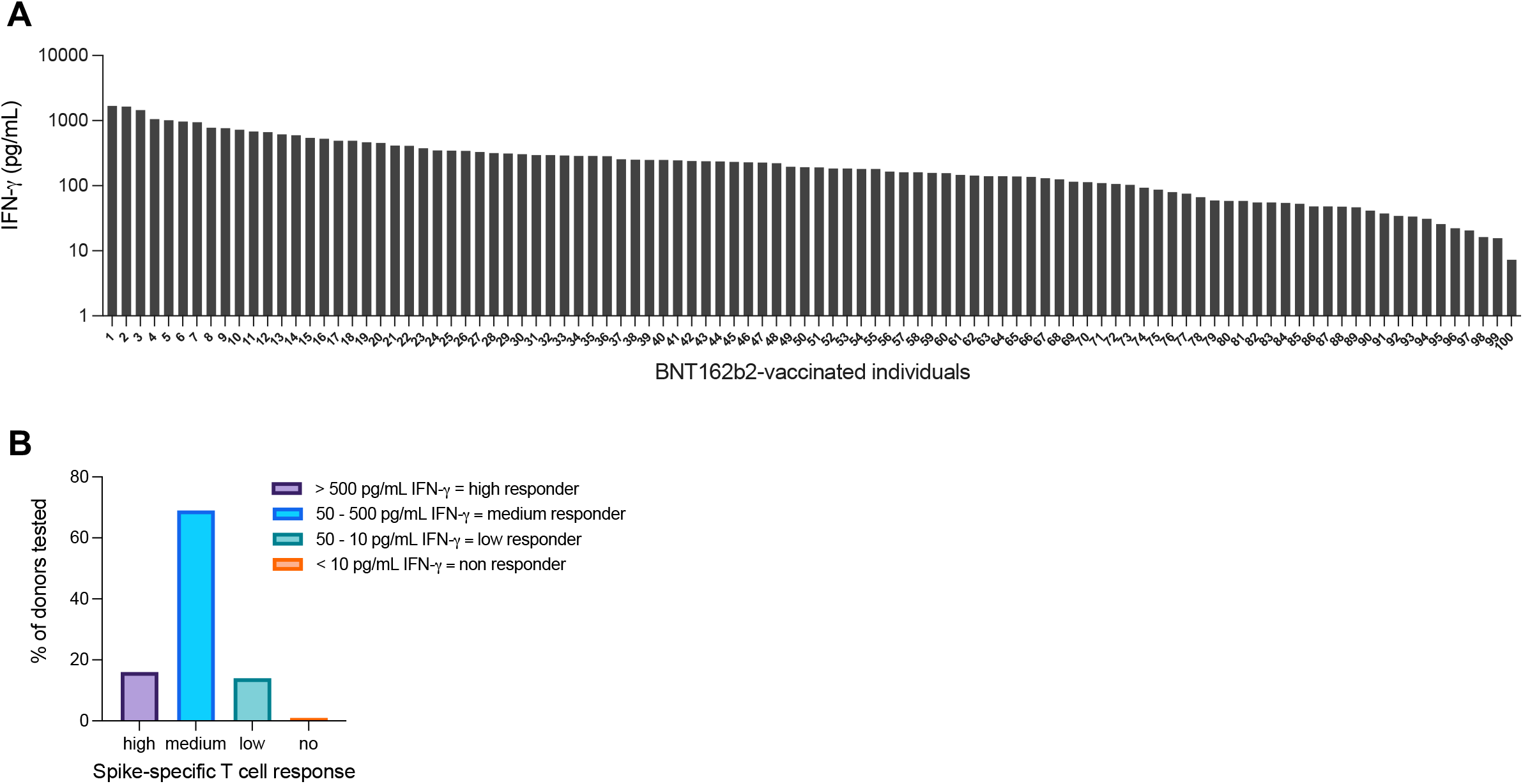
Heterogeneity of the Spike-specific T cell response in BNT162b2 vaccinated donors. **A**, Whole blood of n=100 healthy BNT162b2 vaccinated donors was stimulated with one peptide pool covering the whole Spike protein. IFN-γ release was quantified after overnight stimulation in the plasma. **B**, Donors were divided based on peptide pool reactivity in high, medium, low and non-responder.

**Figure S3:**
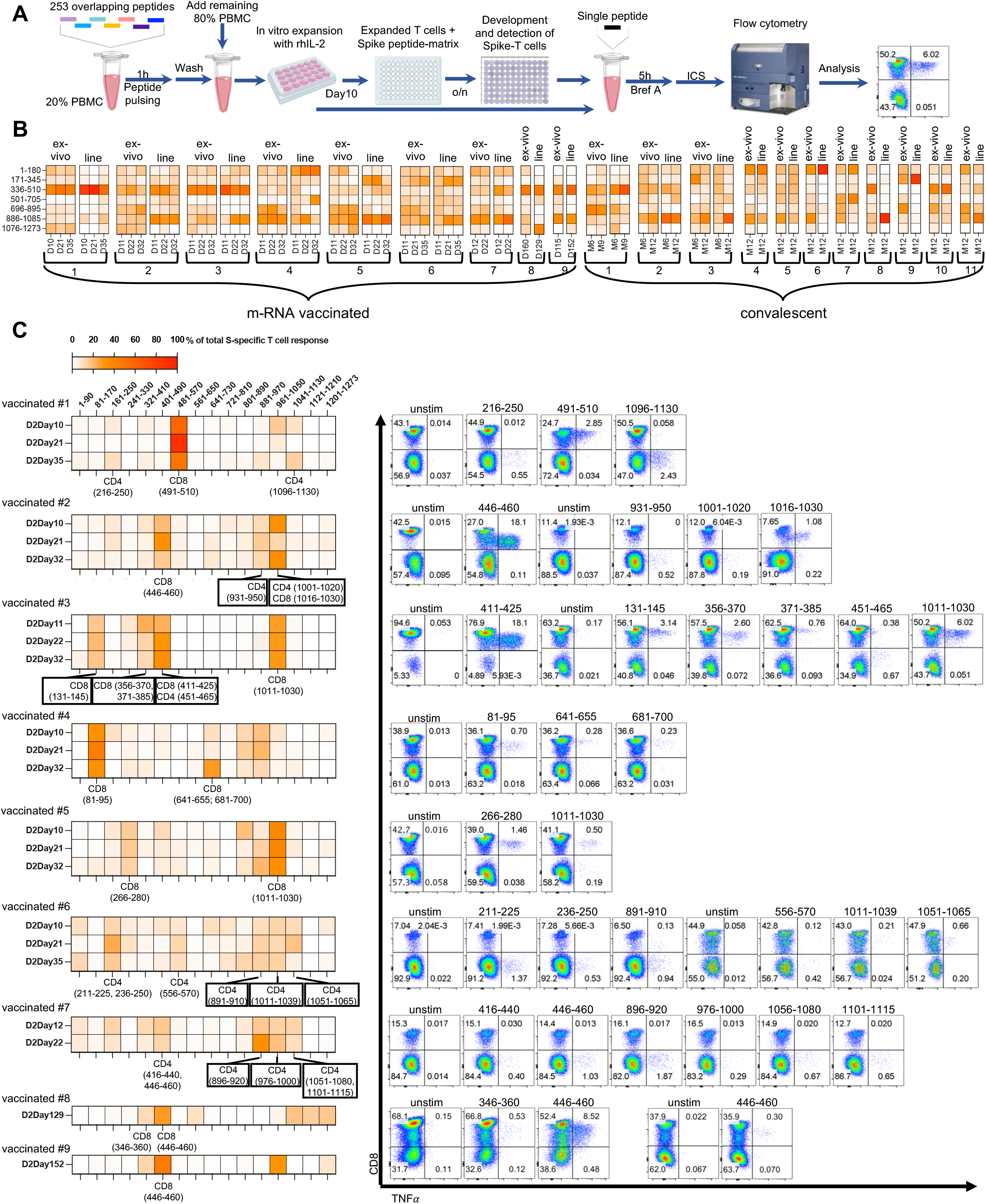
Definition of single-peptide specificities of dominant and subdominant Spike-specific T cells. **A**, Schematic of Spike-specific short-term T cell line expansion procedure, ELISpot peptide-matrix for identification of putative single peptide responses and their confirmation and phenotype characterisation by intracellular cytokine staining and flowcytometry analysis. **B**, Side by side comparison of the response hierarchy to the seven individual Spike-peptide pools directly ex vivo and after in vitro expansion. Heatmap represents the percentage of the response to each peptide pool among the total Spike-response. 9 vaccinated donors and 11 convalescents were analysed at multiple timepoints. **C**, Heatmap of the response of the short-term T cell lines of the 9 vaccinated donors to the numerical pools of the peptide matrix covering 16 distinct regions of the Spike-protein (left). Single-peptide responses were confirmed by ICS (right).

**Figure S4:**
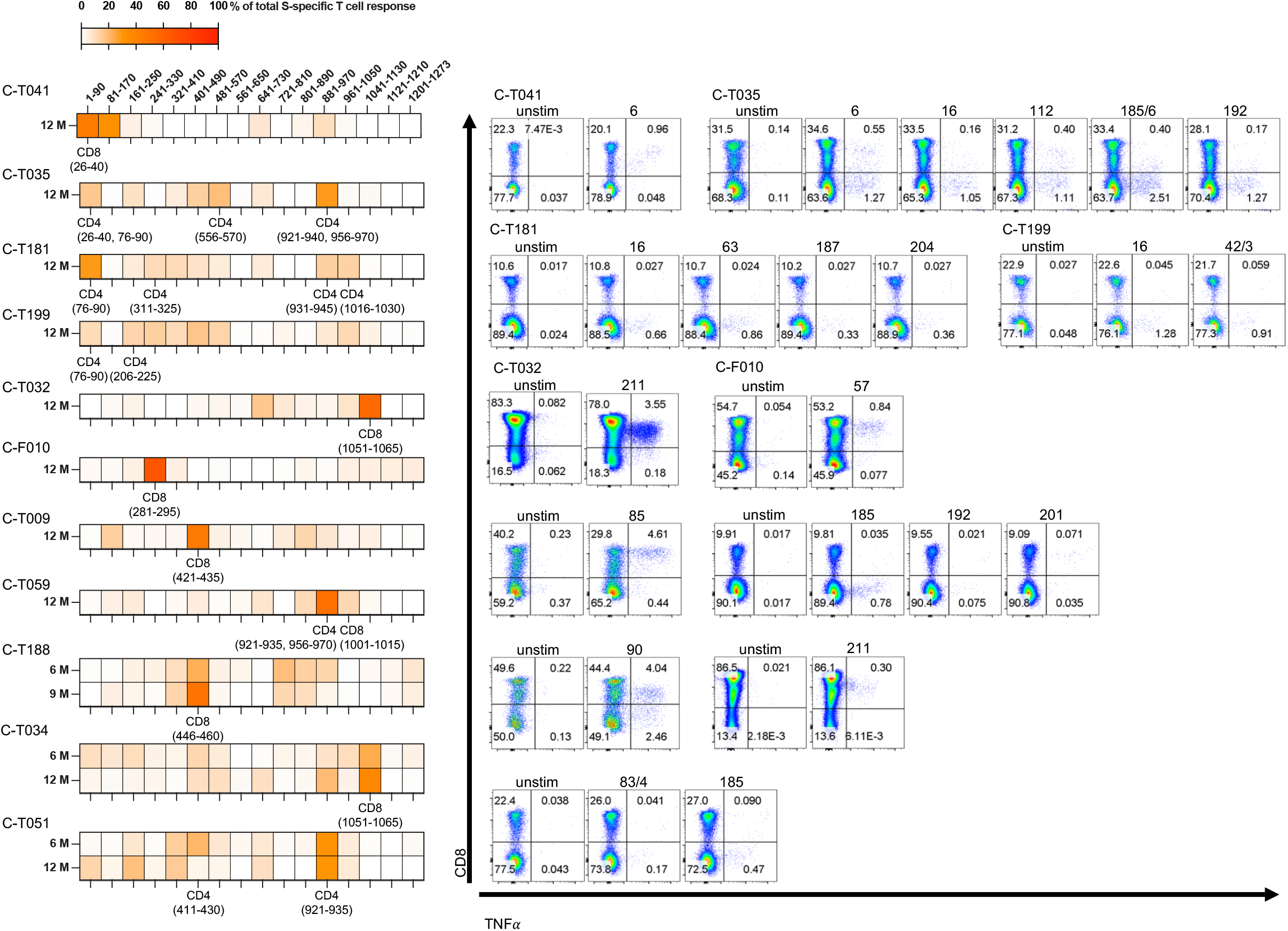
Definition of single-peptide specificities of dominant and subdominant Spike-specific T cells in SARS-CoV-2 convalescent donors. Heatmap of the response of the short-term T cell lines of the 11 convalescent donors to the numerical pools of the peptide matrix covering 16 distinct regions of the Spike-protein (left). Single-peptide responses were confirmed by ICS (right).

**Figure S5:**
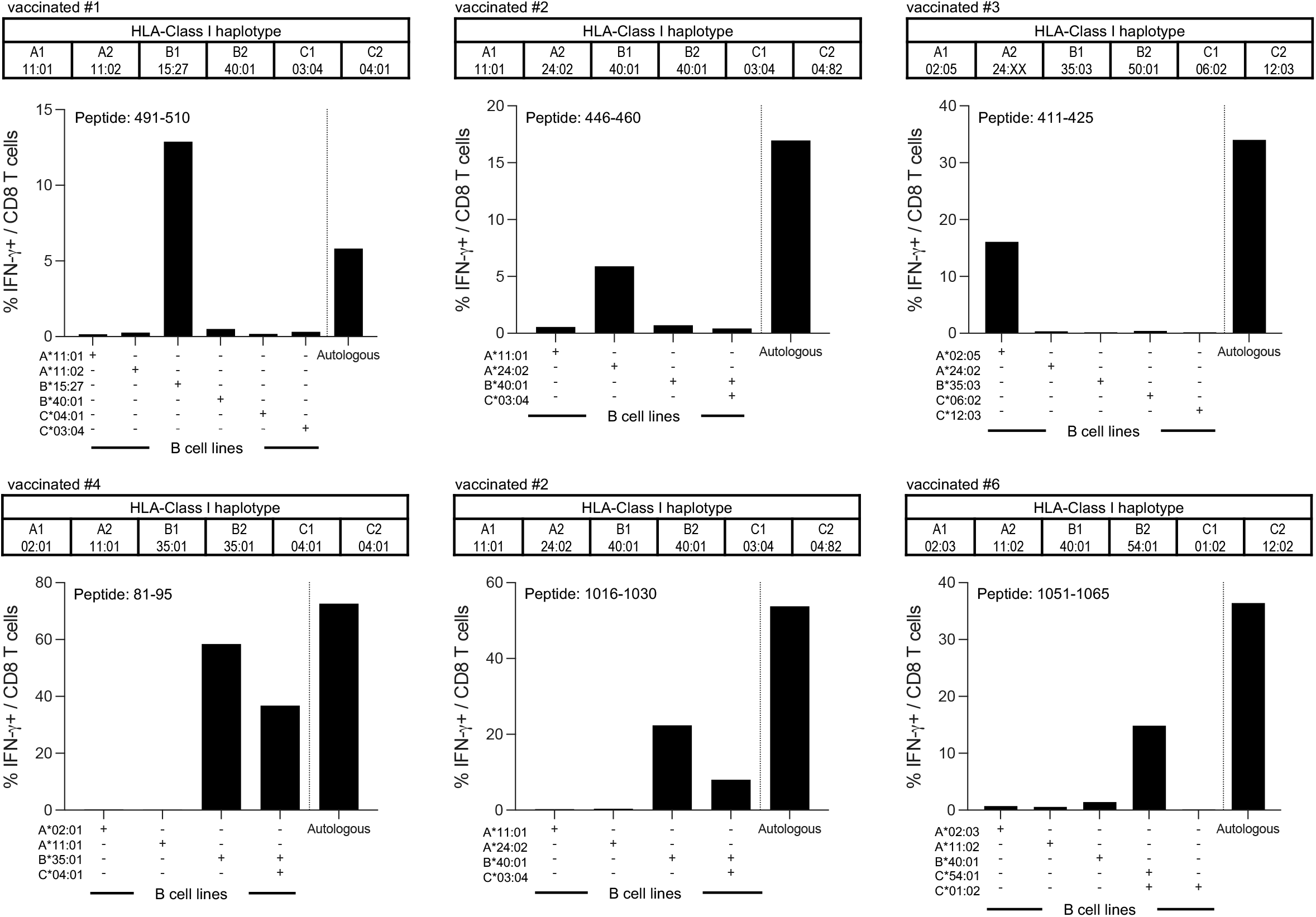
HLA-Class-I restriction of single-peptide specificities of 6 dominant CD8+ Spike-specific T cells. The HLA-class I haplotypes of vaccinated donors 1, 2, 3, 4 are 6 are shown in the tables. HLA-restriction of the indicated peptide-specific T cells from the donors was deduced by co-culturing the T cells with peptide-pulsed EBV-transformed B cell lines that shared the indicated HLA-Class I molecule (+). Activation of the peptide-specific T cells by autologous cells was achieved by the direct addition of the peptide and used as a positive control.

**Figure S6:**
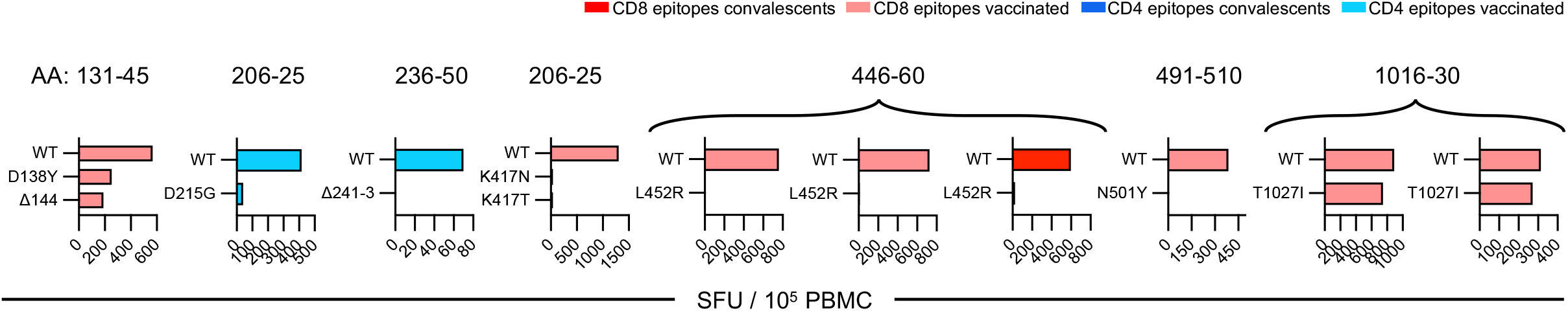
The impact of VOC AA mutations on Spike-specific T cells after in vitro expansion. Expanded T cell lines of the individuals in whom the T cell specificities were defined (and cells were available) were stimulated in parallel with peptides containing either the wildtype or the mutated sequence. Peptide reactivity was analysed by IFN-γ ELISpot assay. Frequency of spot forming cells (SFC)/105 PBMC are shown.

**Figure S7:**
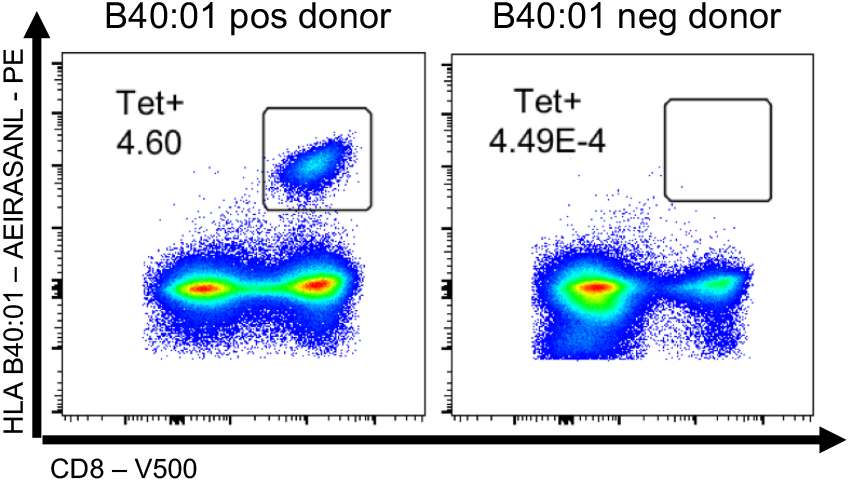
Definition of a CD8 T cell epitope in peptide S1016-30-specific T cells. S1016-30 expanded T cell lines of a HLA-B*40:01+ and a HLA-B*40:01-vaccine recipient were stained with HLA-B*40:01 tetramer containing peptide 1016-24. Dot plots show tetramer staining on CD3+ cells.

**Table S1:**
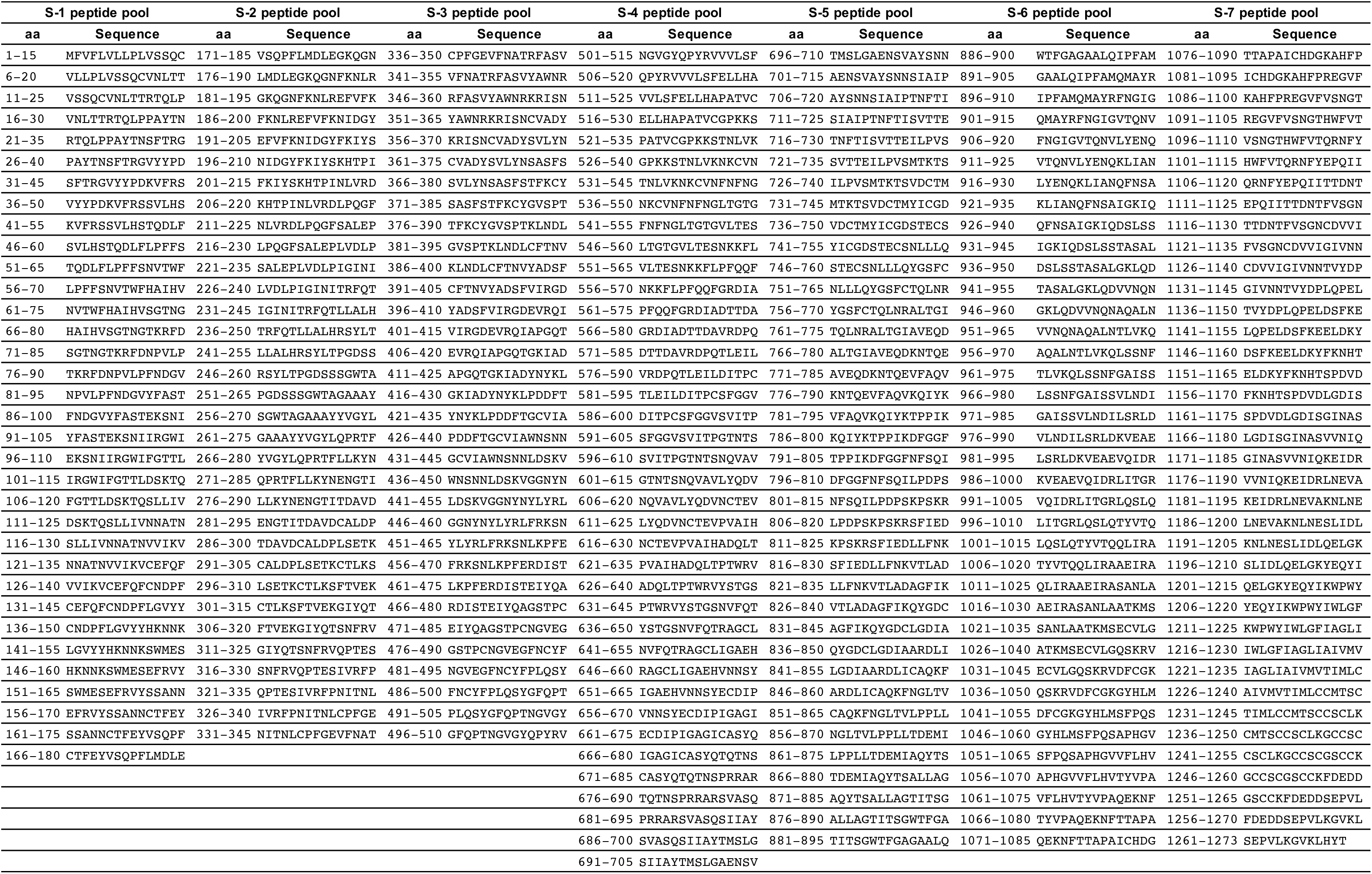
Seven pools of 15-mer overlapping peptides covering the SARS-CoV-2 Spike protein

**Table S2:**
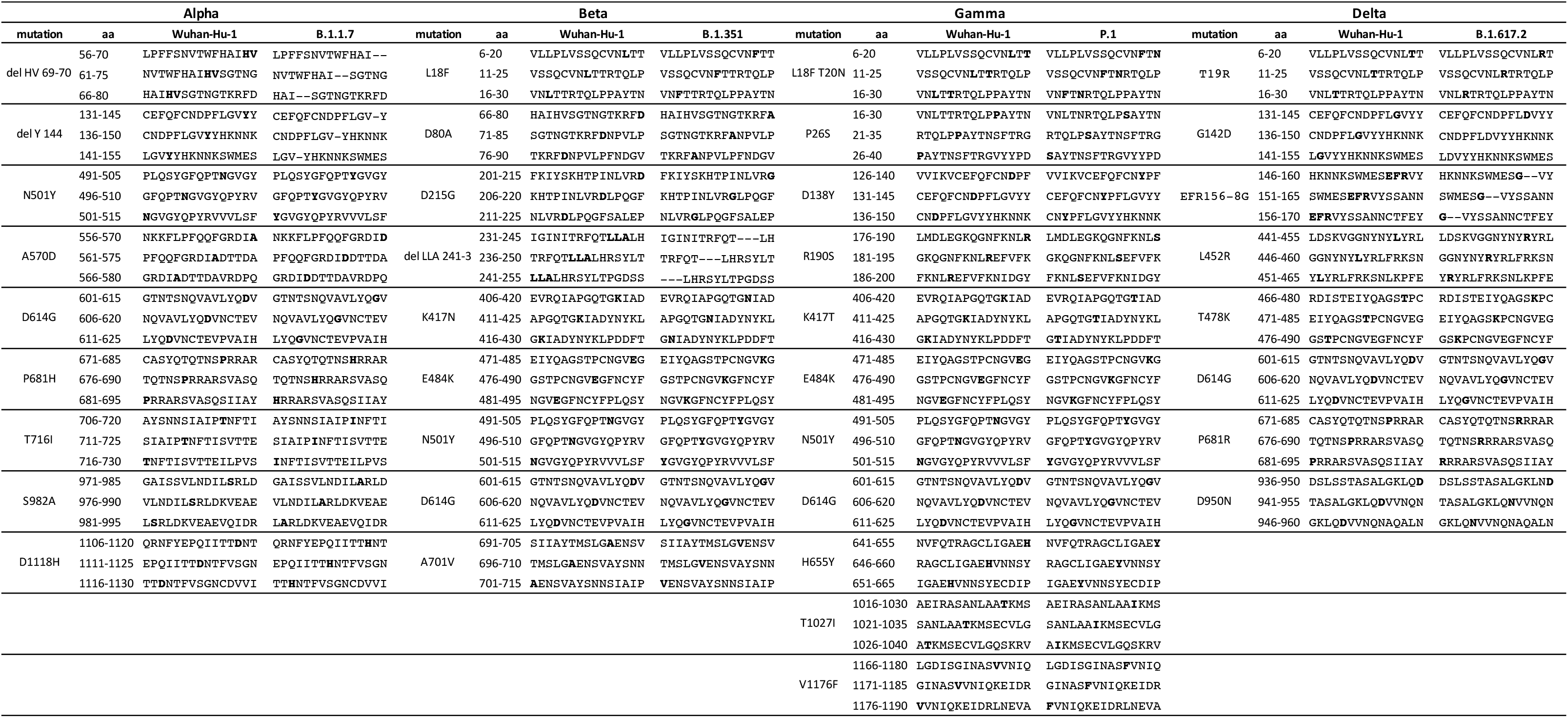
Overlapping 15-mer peptides covering the region of the VOC mutations with and without the amino acid substitutions and deletions.

**Table S3:** Spike mutations in VOCs vs Wuhan-Hu1. Grey highlighted mutations were tested in this manuscript.

|  | Alpha | Beta | Gamma | Delta | Omicron |
| --- | --- | --- | --- | --- | --- |
|  | B.1.1.7 | B.1.351 | P.1 | B.1.617.2 | B.1.1.529 |
| L18F |  | (F) | F |  |  |
| T19R |  |  |  | R |  |
| T20N |  |  | N |  |  |
| P26S |  |  | S |  |  |
| A67V |  |  |  |  | V |
| del 69-70 | del |  |  |  | del |
| D80A |  | A |  |  |  |
| T95I |  |  |  | (I) | I |
| D138Y |  |  | Y |  |  |
| G142D |  |  |  | (D) | D |
| del 143-5 |  |  |  |  | del |
| del 144 | del |  |  |  |  |
| E156G |  |  |  | G |  |
| del 157-8 |  |  |  | del |  |
| R190S |  |  | S |  |  |
| N211I |  |  |  |  | I |
| del 212 |  |  |  |  | del |
| ins EPE 214 |  |  |  |  | ins |
| D215G |  | G |  |  |  |
| del 241-3 |  | del |  |  |  |
| G339D |  |  |  |  | D |
| S371L |  |  |  |  | L |
| S373P |  |  |  |  | P |
| S375F |  |  |  |  | F |
| K417N/T |  | N | T |  | (N) |
| L452R |  |  |  | R |  |
| S477N |  |  |  |  | N |
| T478K |  |  |  | K | K |
| E484A/K |  | K | K |  | A |
| Q493R |  |  |  |  | R |
| G496S |  |  |  |  | S |
| Q498R |  |  |  |  | R |
| N501Y | Y | Y | Y |  | Y |
| Y505H |  |  |  |  | H |
| T547K |  |  |  |  | K |
| A570D | D |  |  |  |  |
| D614G | G | G | G | G | G |
| H655Y |  |  | Y |  | Y |
| N679K |  |  |  |  | K |
| P681R/H | H |  |  | R | H |
| A701V |  | V |  |  | (V) |
| T716I | I |  |  |  |  |
| D796Y |  |  |  |  | Y |
| N856K |  |  |  |  | K |
| D950N |  |  |  | N |  |
| Q954H |  |  |  |  | H |
| N969K |  |  |  |  | K |
| L981F |  |  |  |  | F |
| S982A | A |  |  |  |  |
| T1027I |  |  | I |  |  |
| D1118H | H |  |  |  |  |
| V1176F |  |  | F |  |  |

## Notes

### Competing Interest Statement

The authors have declared no competing interest.

